# Intracellular flow cytometry staining of antibody-secreting cells using phycoerythrin-conjugated antibodies: pitfalls and solutions

**DOI:** 10.1101/2022.01.10.475671

**Authors:** Patrick Renner, Michael Crone, Matthew Kornas, KimAnh T. Pioli, Peter D. Pioli

**Affiliations:** Center for Immunobiology, 1000 Oakland Drive, Kalamazoo, MI, 49008 United States; Department of Investigative Medicine, 1000 Oakland Drive, Kalamazoo, MI, 49008 United States; Western Michigan University Homer Stryker M.D. School of Medicine, 1000 Oakland Drive, Kalamazoo, MI, 49008 United States

**Keywords:** antibody-secreting cell, plasma cell, plasmablast, phycoerythrin, intracellular flow cytometry

## Abstract

Antibody-secreting cells are terminally differentiated B cells that play a critical role in humoral immunity through immunoglobulin secretion along with possessing the potential to be long-lived. It is now appreciated that antibody-secreting cells regulate multiple aspects of biology through the secretion of various cytokines. In this regard, intracellular flow cytometry is a key tool used to assess the presence of intracellular proteins such as cytokines and transcription factors. Here, we showed that the use of phycoerythrin-containing antibody conjugates led to a false interpretation of antibody-secreting cell intracellular protein expression compared to other cell types. This was mainly due to the inappropriate retention of these antibodies specifically within antibody-secreting cells. Furthermore, we demonstrated how to reduce this retention which allowed for a more accurate comparison of intracellular protein expression between antibody-secreting cells and other cell types such as B lymphocytes. Using this methodology, our data revealed that spleen antibody-secreting cells expressed Toll-like receptor 7 as well as the pro-form of the inflammatory cytokine interleukin-1β.

## 1. Introduction

Humoral, or antibody (Ab)-mediated, immunity plays a critical role in host defense (Amanna et al., 2007). Through a variety of mechanisms, the activation of B cells can lead to their terminal differentiation into antibody-secreting cells (ASCs) which include relatively short-lived plasmablasts (PBs) as well as more mature plasma cells (PCs) (Fairfax et al., 2008; Gaudette and Allman, 2021; Manakkat Vijay and Singh, 2021; Nutt et al., 2015). In general, ASCs are viewed as critical effectors of humoral immunity through their potential to be long-lived (Landsverk et al., 2017; Manz et al., 1998) as well as their continuous production of Abs (Slifka et al., 1998).

It is now appreciated that ASCs have the ability to influence host biology well beyond the direct effects of Ab synthesis (Ma et al., 2020; McGettigan and Debes, 2021; Pioli, 2019). Given that ASCs are programmed for protein production and secretion (Shaffer et al., 2004; Zhu et al., 2019), it is not a surprise that many of the functions they exert are mediated by cytokine production. For example, ASCs utilize interleukin (IL)-10 to suppress the inflammatory response in the context of autoimmunity (Matsumoto et al., 2014; Rojas et al., 2019). However, ASCs can also facilitate the immune response through the production of factors including IL-6 (Chavele et al., 2015) as well as IL-3 (Chin et al., 2019) and granulocyte-macrophage colony-stimulating factor (GM-CSF) (Rauch et al., 2012; Weber et al., 2014).

Multiple strategies currently exist which allow for the evaluation of ASC cytokine production. For instance, ASCs can be isolated via fluorescence-activated cell sorting, stimulated *in vitro* and have culture supernatants directly probed for selected cytokines (Lino et al., 2018). While informative, this approach requires knowing the correct stimulus, or stimuli, and does not necessarily provide insight into the cytokines that ASCs are actively producing while residing within a particular tissue. As such, this type of analysis has been paired with *in vivo* techniques that utilize fluorescent reporter mice such as those specific for IL-10 (Lino et al., 2018; Matsumoto et al., 2014; Meng et al., 2019; Shen et al., 2014; Suzuki-Yamazaki et al., 2017). However, since genetically engineered reporter mice are not readily available for most cytokines, intracellular flow cytometry (ICFC) is also extensively performed to assess ASC cytokine production (Chin et al., 2019; Rauch et al., 2012; Weber et al., 2014). Indeed, this approach was recently used to show that plasmablasts produced IL-3 and GM-CSF in the context of malaria-inducing parasitic infections (Chin et al., 2019).

Within the field of flow cytometry, phycoerythrin (PE)-conjugated Abs are known to be extremely “bright” and provide large signal-to-noise ratios thus facilitating their use to detect rare protein targets (Batard et al., 2002). However, an under-appreciated study has previously shown that ASCs non-specifically retain PE during ICFC (Kim and Kim, 2013). This artifact pertains to *ex vivo* stained ASCs and well as those derived *in vitro*. Very recently, non-specific PE retention was observed when ICFC was performed on day 3 cultures from lipopolysaccharide (LPS)-stimulated B cells (Bohacova et al., 2021), a timepoint known to generate a significant proportion of ASCs. Quite clearly, this retention phenomenon is a confounding factor in the analysis of ASC cytokine production and evaluation with ICFC.

In this study, we showed that the complication of non-specific PE retention by ASCs also pertained to PE-containing tandem fluorochromes such as PE/Cy7. Interestingly enough, the number of centrifugation steps post-fixation directly modulated the extent of this retention. Provided here is a standard approach that reduces PE retention thus allowing the use of PE- and PE/Cy7-conjugates Abs to perform ICFC on ASCs. Via this methodology, we demonstrated that spleen (SPL) ASCs possessed increased amounts of Pro-IL-1β compared to B220^+^ CD138^-^ B cells (BCs) from the SPL. Furthermore, we detected TLR7 protein expression by ASCs thus adding to the growing list of immunomodulatory receptors that these cells possess.

## 2. Methods

### 2.1. Mice

B6.Cg-Tg(Prdm1-EYFP)1Mnz/J (Stock #: 008828) and wildtype C57BL/6 mice were purchased from the Jackson Laboratory and used to establish a breeding colony within the vivarium of the Western Michigan University Homer Stryker M.D. School of Medicine (WMed). Wildtype mice were used for all experiments. Both females and males were used between 12-32 weeks of age for experiments in Figures 1–4 and Supplementary Figure 1. 25-weeks old males were specifically used in Figure 5 and Supplementary Figure 2 to correct for any potential age- or sex-based effects on target protein expression. All experiments were performed with the approval of the WMed Institutional Animal Care and Use Committee (IACUC).

### 2.2. Isolation of spleen tissue

SPLs were processed and collected in calcium and magnesium-free 1x phosphate buffered saline (PBS). Organs were crushed between the frosted ends of 2 slides and cell suspensions were centrifuged for 5 minutes at 4 °C and 600g. Red blood cells were lysed by resuspending (RSS) cells in 3 mL of 1x red blood cell lysis buffer on ice for ~3 minutes. Lysis was stopped with the addition of 7 mL of 1x PBS. Cell suspensions were counted via a hemocytometer using Trypan Blue to exclude dead cells and subsequently passed through 70-μm filters, centrifuged as above and RSS in 1x PBS + 0.1% bovine serum albumin (BSA) at a concentration of 2 x 10^7^ cells/mL before use.

### 2.3. Cell surface immunostaining for flow cytometry

All cell surface staining procedures were performed in 1x PBS + 0.1% BSA using ~5 x 10^6^ cells per stain. Samples were labeled with a CD16/32 blocking Ab to eliminate non-specific binding of Abs to cells via Fc receptors. All Abs utilized are listed in the Supplementary Table 1. Cells were incubated on ice for 30 minutes with the appropriate Abs. Unbound Abs were washed away with 3 mL of 1x PBS + 0.1% BSA followed by centrifugation for 5 minutes at 4 °C and 600g. Supernatants were decanted. In certain instances, eBioscience Fixable Viability (Live-Dead) Dye eFluor 780 (Thermo Fisher Scientific, Catalog # 65-0865-14) was added to samples to assess dead cell content. The stock solution was diluted 1:250 and 10 uL was added to ~5 x 10^6^ cells per stain. Live-Dead stain was added concurrent with surface staining Abs.

### 2.4. Paraformaldehyde and saponin immunostaining for intracellular flow cytometry

Cells were surface stained as described above. Afterwards, cell pellets were RSS in 1 mL of 4% paraformaldehyde (PFA) and incubated at room temperature (RT) for 20 minutes in the dark. Subsequently, 2 mL of 1x PBS + 0.1% BSA + 0.1% Saponin (Sap) were added and cells centrifuged for 5 minutes at 4 °C and 600g. Supernatants were decanted and the previous wash step was repeated with 3 mL of 1x PBS + 0.1% BSA + 0.1% Sap for a total of 2 washes. Supernatants were discarded and cells were RSS in residual buffer (~150 μL). Unlabeled mouse IgG was added as a blocking reagent and cells were incubated at RT for 20 minutes in the dark. Subsequently, the appropriate Abs were added and cells were incubated at RT for 30 minutes in the dark. Afterwards, 3 mL of 1x PBS + 0.1% BSA + 0.1% Sap were added and cells centrifuged for 5 minutes at 4 °C and 600g. Supernatants were decanted and cells were washed with 3 mL of 1x PBS + 0.1% BSA. Supernatants were discarded and cell pellets were RSS in an appropriate volume of 1x PBS + 0.4% BSA + 2 mM EDTA for flow cytometric analysis. Before analysis, cells were strained through a 35 μM filter mesh. Any deviation from wash step constituents or sequence are noted in the Results section.

### 2.5. eBioscience Foxp3/Transcription Factor Staining Buffer Set immunostaining for intracellular flow cytometry

For this procedure, ICFC was performed with the eBioscience Foxp3/Transcription Factor Staining Buffer Set (Thermo Fisher Scientific, Catalog # 00-5523-00). Cells were surface stained as described above. Afterwards, cell pellets were RSS in 1 mL of Buffer 1 and incubated at RT for 20 minutes in the dark. Subsequently, 2 mL of Buffer 2 were added and cells centrifuged for 5 minutes at 4 °C and 600g. Supernatants were decanted and the previous wash step was repeated with 2 mL of Buffer 2 for a total of 2 washes. Supernatants were discarded and cells were RSS in residual buffer (~150 μL). Unlabeled mouse IgG was added as a blocking reagent and cells were incubated at RT for 20 minutes in the dark. Subsequently, the appropriate Abs were added and cells were incubated at RT for 30 minutes in the dark. Afterwards, 2 mL of Buffer 2 were added and cells centrifuged for 5 minutes at 4 °C and 600g. Supernatants were decanted and cells were washed with 3 mL of 1x PBS + 0.1% BSA. Supernatants were discarded and cell pellets were RSS in an appropriate volume of 1x PBS + 0.4% BSA + 2 mM EDTA for flow cytometric analysis. Before analysis, cells were strained through a 35 μM filter mesh. Any deviation from wash step constituents or sequence are noted in the Results section.

### 2.6. Flow cytometry

Flow cytometry was performed on a Fortessa (BD Biosciences) located in the Flow Cytometry and Imaging Core at WMed. Data were analyzed using FlowJo (v10) software. Total cells were gated using side scatter area (SSC-A) versus forward scatter (FSC-A) area. Singlets were identified using sequential gating of FSC-width (W) versus FSC-A, SSC-W versus SSC-A and FSC-height (H) versus FSC-A.

### 2.7. Quantification and statistical analysis

Technical and biological replicates are indicated in the Figure Legends.

## 3. Results

### 3.1. Antibody-secreting cells preferentially retain phycoerythrin-conjugated antibodies

ICFC of cytokines has become a critical tool used to evaluate ASC diversification and function. Using conventional fixation (i.e. PFA) and permeabilization (i.e. Sap) techniques, it has been reported that ASCs non-specifically retain PE-conjugated Abs thus making them unsuitable for ICFC (Kim and Kim, 2013). This was shown to be the case for *ex vivo* stained ASCs as well as those generated *in vitro* following B lymphocyte stimulation (Kim and Kim, 2013).

To better characterize this phenomenon, we evaluated the ability of B220^+/-^ CD138^HI^ CD90.2^-^ IgD^-^ ASCs (**Figure 1A**) from the SPL to retain a variety of non-specific isotype control Abs conjugated to allophycocyanin (APC), PE and PE/Cy7. As a point of comparison, we also examined this capacity in SPL B220^+^ CD138^-^ BCs (**Figure 1A**). For these experiments, we utilized hamster (ham) IgG, rat (r) IgG1, rIgG2a and rIgG2b isotypes. Using flow cytometry, retention was measured as a function of geometric mean fluorescence intensity (gMFI). In this regard, increased gMFIs were indicative of increased retention (**Supplementary Figures 1A-F**).

**Figure 1.**
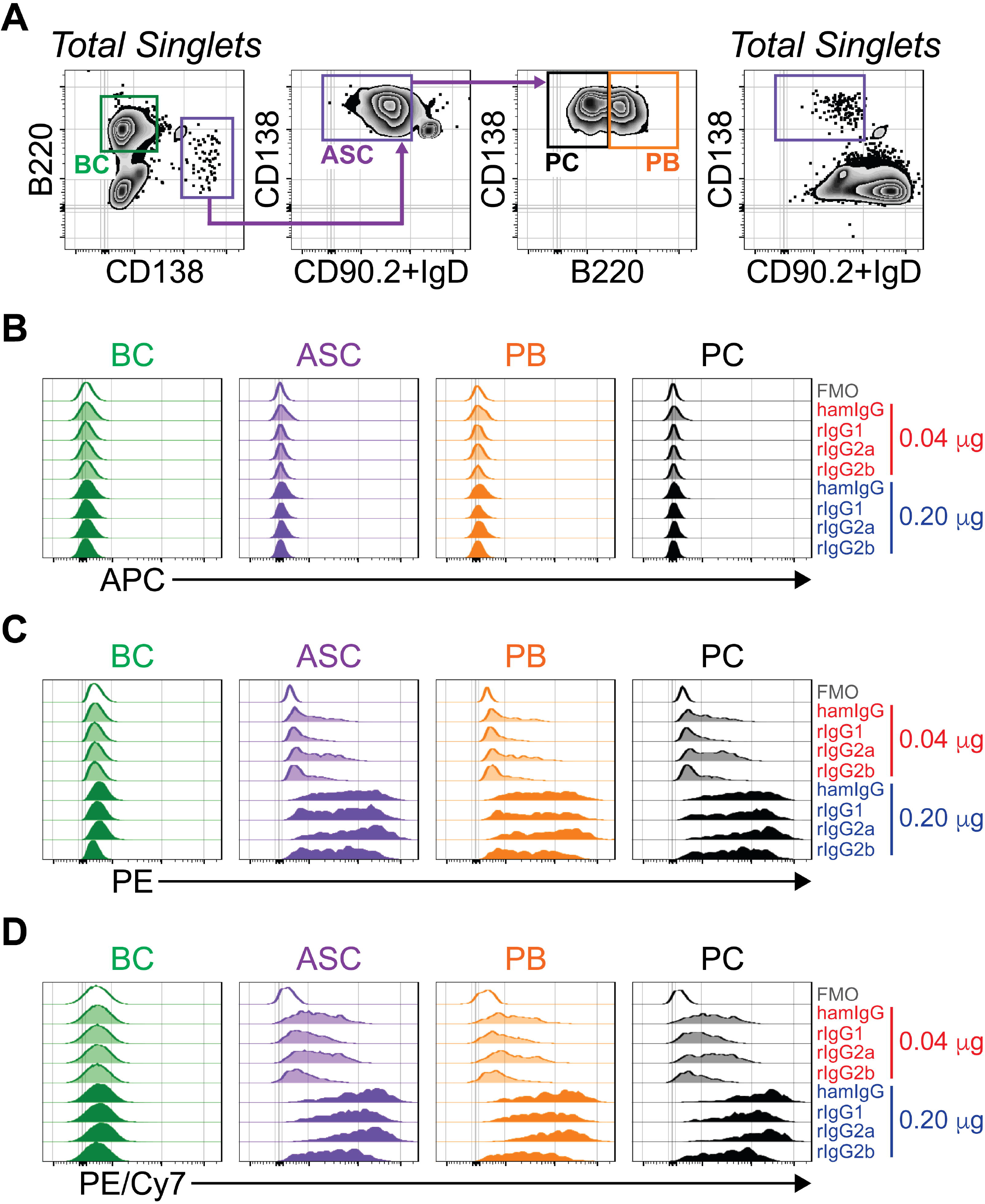
ASCs non-specifically retain PE- and PE/Cy7-conjugated Abs. **(A)** Representative flow cytometry plots demonstrating gating of SPL B220^+^ CD138^-^ BCs and B220^+/-^ CD138^HI^ CD90.2^-^ IgD^-^ ASCs. Total ASCs were further subdivided into B220^+^ PB and B220^-^ PC. Cells are pre-gated on total singlets. CD138 versus CD90.2+IgD staining of total singlets in shown for reference. **(B-D)** Flow cytometry overlay histograms showing fluorescence intensity for **(B)** APC, **(C)** PE and **(D)** PE/Cy7. FMO and isotype control Ab (0.04 and 0.020 μg) stains are shown for SPL BCs, total ASCs, PBs and PCs. Data are representative of 3 individual experiments and gMFIs are summarized in **Supplementary Fig. 1**.

Abs conjugated to APC produced very little background when compared to fluorescence minus one (FMO) controls (**Figure 1B and Supplementary Figure 1A**). This was independent of concentration and observed in both BCs and ASCs. However, the examination of PE-conjugated isotype Abs painted a different picture. While BC and ASC FMO controls possessed similar amounts of PE-mediated background autofluorescence (**Figure 1C and Supplementary Figure 1C**), the use of low amounts (0.04 μg) of PE-conjugated isotype Abs resulted in low but evident levels of retention in ASCs relative to BCs (**Figure 1C and Supplementary Figure 1C**). The use of increased Ab amounts (0.20 μg) led to further retention by ASCs which was apparent for all isotypes (**Figure 1C and Supplementary Figure 1C**). To determine if these observations could be more generalized, we examined retention of the tandem dye PE/Cy7 (**Figure 1D and Supplementary Figure 1E**). Similar to PE, PE/Cy7 retention by ASCs was evident in a concentration-dependent manner regardless of isotype (**Figure 1D and Supplementary Figure 1E**).

Additionally, we subdivided ASCs (**Figure 1A**) into short-lived plasmablasts (PBs) and more mature, potentially long-lived, plasma cells (PCs) which are B220^+^ and B220^-^, respectively. Analysis of APC-, PE- and PE/Cy7-conjugated isotype Ab retention demonstrated a high degree of similarity between PBs and PCs (**Figures 1B-D and Supplementary Figures 1B, 1D and 1F**). These data indicated that maturation status of ASCs (PBs vs PCs) did not explicitly alter the capacity to retain PE- and PE/Cy7-conjugated isotype Abs. Furthermore, these results suggested that potential contamination with non-ASCs that possess B220 expression (e.g. plasmacytoid dendritic cells) did not heavily influence the data. In summary, ASCs displayed retention of PE-based fluorochromes that was independent of Ab isotype and species but was augmented by increased Ab amounts.

### 3.2. The number of washes following paraformaldehyde fixation modulates antibody-secreting cell retention of phycoerythrin-conjugated antibodies

In our staining protocol, we utilized one post-stain wash that included Sap (**Figure 2A**, **brown**). This was then followed by a wash with PBS + 0.1% BSA. Therefore, it is possible that additional Sap washes could abrogate PE retention. To test this, we performed experiments in which cells stained with rIgG2a-PE (0.20 μg) received 1, 2 or 3 post-stain Sap washes followed by a terminal PBS + 0.1% BSA wash (**Figures 2B-D**). The number of post-stain Sap washes had essentially no effect on BC background PE staining (**Figures 2B-C**). The addition of a 2^nd^ or even 3^rd^ wash had only minor effect on PE retention by ASCs compared to cells that received a single wash (**Figure 2D**). The ASC / BC PE gMFI ratio (**Figure 2D**) ranged from ~14-33 regardless of the number of post-stain Sap washes. These data confirmed previous observations from studies in which multiple post-stain washes containing permeabilizing agents were ineffectively used to remove PE retention by ASCs (Bohacova et al., 2021; Kim and Kim, 2013).

**Figure 2.**
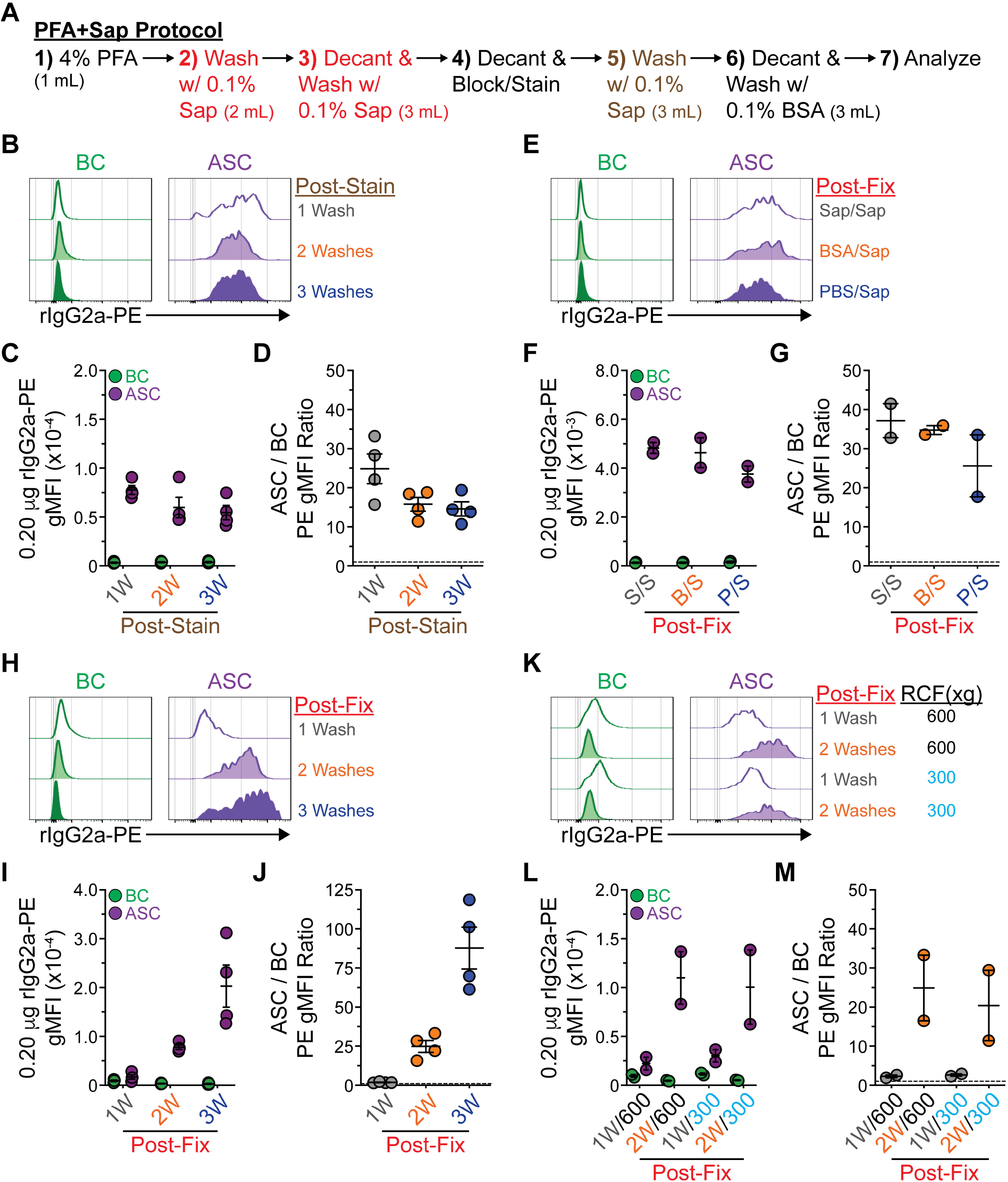
The number of post-fixation washes modulates ASC retention of PE- and PE/Cy7-conjugated Abs. **(A)** Schematic depicting PFA+Sap staining protocol. **(B)** Flow cytometry overlay histograms showing rIgG2a-PE fluorescence intensity for SPL BCs and ASCs using 1, 2 or 3 post-stain Sap washes. **(C)** rIgG2a-PE (0.20 μg) gMFIs of SPL BCs and ASCs washed 1, 2 or 3 times with Sap following staining. **(D)** Ratio of SPL ASC / BC rIgG2a-PE (0.20 μg) gMFIs from cells washed 1, 2 or 3 times with Sap following staining. Horizontal dashed line represents a ratio of 1. **(E)** Flow cytometry overlay histograms showing rIgG2a-PE fluorescence intensity for SPL BCs and ASCs using Sap/Sap, BSA/Sap or PBS/Sap 2-step washes post-fixation. **(F)** rIgG2a-PE (0.20 μg) gMFIs of SPL BCs and ASCs washed with Sap/Sap (S/S), BSA/Sap (B/S) or PBS/Sap (P/S). **(G)** Ratio of SPL ASC / BC rIgG2a-PE (0.20 μg) gMFIs from cells washed with S/S, B/S or P/S. Horizontal dashed line represents a ratio of 1. **(H)** Flow cytometry overlay histograms showing rIgG2a-PE fluorescence intensity for SPL BCs and ASCs using 1, 2 or 3 post-fixation Sap washes. **(I)** rIgG2a-PE (0.20 μg) gMFIs of SPL BCs and ASCs washed 1, 2 or 3 times with Sap following fixation. **(J)** Ratio of SPL ASC / BC rIgG2a-PE (0.20 μg) gMFIs from cells washed 1, 2 or 3 times with Sap following fixation. Horizontal dashed line represents a ratio of 1. **(K)** Flow cytometry overlay histograms showing rIgG2a-PE fluorescence intensity for SPL BCs and ASCs using 1 or 2 post-fixation Sap washes at 600g or 300g. **(L)** rIgG2a-PE (0.20 μg) gMFIs of SPL BCs and ASCs washed 1 or 2 times at 600g or 300g with Sap following fixation. **(M)** Ratio of SPL ASC / BC rIgG2a-PE (0.20 μg) gMFIs from cells washed 1 or 2 times at 600g or 300g with Sap following fixation. Horizontal dashed line represents a ratio of 1. **(B-D, H-J)** Data derived from 4 separate experiments. **(E-G, K-M)** Data derived from 2-4 separate experiments. **(C-D, F-G, I-J, L-M)** Symbols represent individual mice. Horizontal lines represent mean ± SEM.

The experiments presented in **Figure 1** utilized 2 post-fixation washes with 0.1% Sap as part of a PBS + 0.1% BSA solution (**Figure 2A, red**). While it is commonly appreciated that fixation method can alter immunostaining properties of cells (Matsuda et al., 2011), it is not readily known how the steps used to remove a fixative can alter immunostaining. To test this, we kept the 2^nd^ post-fixation wash constant and performed the 1^st^ wash with PBS + 0.1% BSA + 0.1% Sap (Sap/Sap), PBS + 0.1% BSA (BSA/Sap) or PBS (PBS/Sap). Subsequently, we analyzed retention of rIgG2a-PE (0.20 μg) by BCs and ASCs (**Figure 2E**). Regardless of 1^st^ wash composition, ASCs still retained rIgG2a-PE as shown by heightened PE gMFIs when compared to BCs (**Figures 2E-F**). Ultimately, rIgG2a-PE background was increased ~30x in ASCs (**Figure 2G**).

Since the number of post-stain Sap washes as well as post-fixation wash buffer composition did not appear to play a major role, we considered “physical” forces involved in ICFC that may augment ASC retention of PE. ASCs are complex cells whose cytoplasm is dominated by an extensive endoplasmic reticulum and Golgi network (Ribatti, 2017), which is quite a contrast to naive BCs. As such, the act of centrifugation and the forces imparted on BCs and ASCs may be different and thus lead to ASC-specific PE retention. Along these lines, variations in centrifugation protocols can have dramatic effects on recovery as well as viability of mechanically sensitive cells (Katkov and Mazur, 1999; Kim et al., 2009). To test this hypothesis, we repeated the above retention experiments using either 1, 2 or 3 post-fixation washes with PBS + 0.1% BSA + 0.1% Sap (**Figure 2H**). Increasing the number of washes slightly decreased the already low levels of rIgG2a-PE staining in BCs (**Figures 2H-I**). Amazingly, rIgG2a-PE (0.20 μg) retention by ASCs was dramatically enhanced as more washes and subsequent centrifugation steps were performed (**Figures 2H-I**). When compared to BCs, the rIgG2a-PE staining of ASCs was near equivalence at 1 wash. At 2 washes, the ratio of ASC / BC PE gMFI was ~25 (**Figure 2J**) similar to what was previously shown in **Figure 2D**. Finally, the addition of a 3^rd^ wash dramatically increased this ratio to >93, on average (**Figure 2J**). To further assess the impact of “force” effects on PE retention, we performed experiments in which cells received either 1 or 2 post-fixation washes using a relative centrifugal force (RCF) of 600g or 300g (**Figure 2K**). If total RCF was a determining factor, ASCs which received single or multiple 300g centrifugation steps would display less retention than their 600g counterparts. Somewhat surprisingly, the level of PE retention observed in ASCs correlated with the number of centrifugation steps rather than the amount of RCF imparted on cells (**Figure 2K**). This was readily apparent when we quantified the gMFI of PE retention (**Figure 2L**) as well as the ratio of ASC / BC PE retention (**Figure 2M**). In both instances, ASCs washed twice routinely demonstrated increased levels of PE staining regardless of RCF. Taken in total, these results indicated that the physical act of washing cells post-PFA fixation was the most critical driver in ASC PE retention.

### 3.3. The number of post-fixation washes modulates antibody-secreting cell retention of phycoerythrin-conjugated antibodies when using the eBioscience Foxp3/Transcription Factor Staining Buffer Set

The experiments using PFA+Sap (**Figure 2**) demonstrated that the number of post-fixation wash steps significantly impacted the retention of PE by ASCs. However, it is unknown if this phenomenon is buffer specific or could be observed using alternative buffer sets. To test this, we examined the effects of centrifugation using the eBioscience Foxp3/Transcription Factor Staining Buffer Set (**Figure 3A**). We chose this buffer set as it has been previously used in the context of ICFC and demonstrated PE-retention in day 3 cultures from LPS-stimulated B cells (Bohacova et al., 2021). Notably, this timepoint would be expected to contain a significant proportion of ASCs.

**Figure 3.**
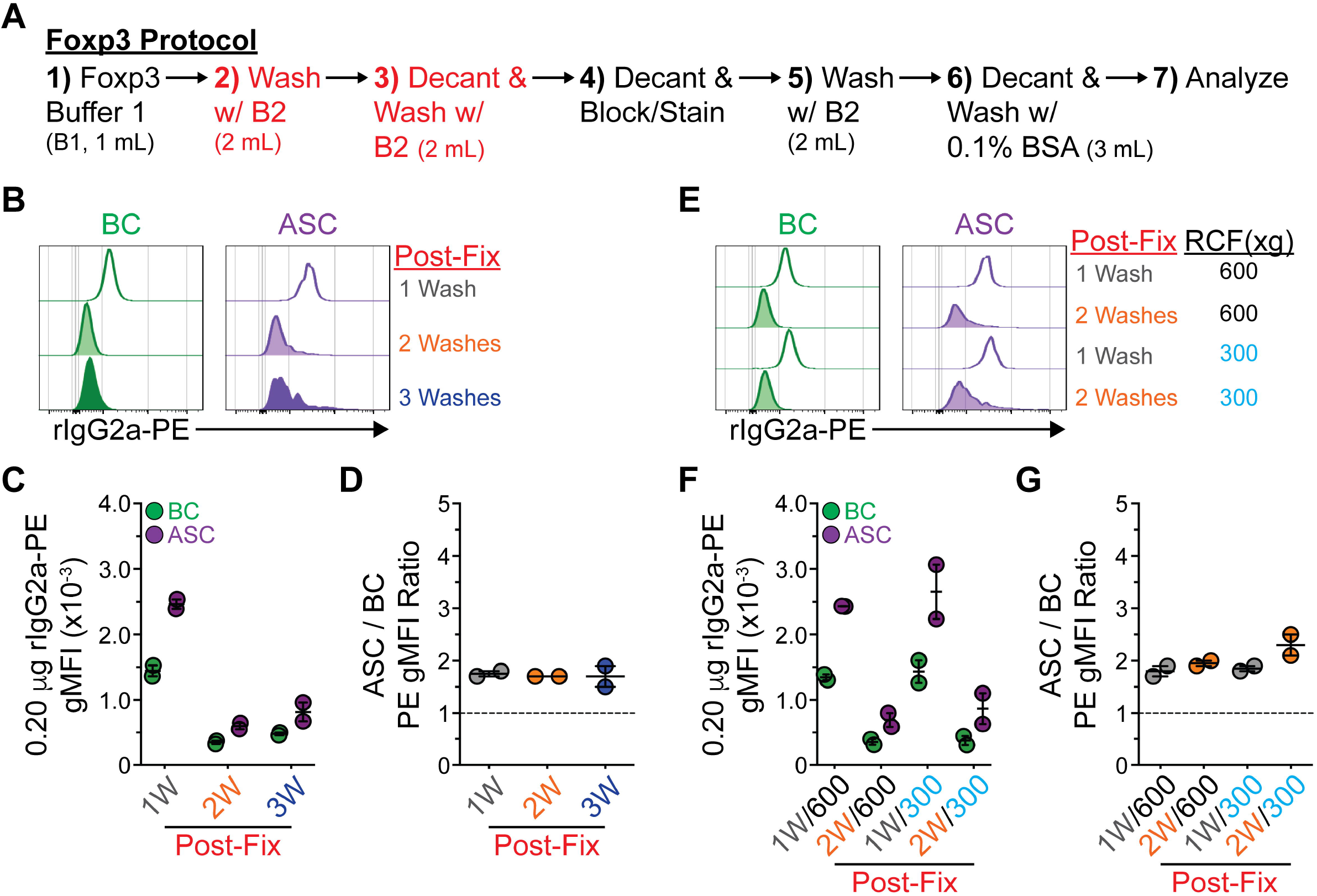
The number of post-fixation washes modulates ASC retention of PE-conjugated Abs when using the eBioscience Foxp3/Transcription Factor Staining Buffer Set. **(A)** Schematic depicting Foxp3 staining protocol. **(B)** Flow cytometry overlay histograms showing rIgG2a-PE fluorescence intensity for SPL BCs and ASCs using 1, 2 or 3 post-fixation Foxp3 B2 washes. **(C)** rIgG2a-PE (0.20 μg) gMFIs of SPL BCs and ASCs washed 1, 2 or 3 times with Foxp3 B2 following fixation. **(D)** Ratio of SPL ASC / BC rIgG2a-PE (0.20 μg) gMFIs from cells washed 1, 2 or 3 times with Foxp3 B2 following fixation. Horizontal dashed line represents a ratio of 1. **(E)** Flow cytometry overlay histograms showing rIgG2a-PE fluorescence intensity for SPL BCs and ASCs using 1 or 2 post-fixation Foxp3 B2 washes at 600g or 300g. **(F)** rIgG2a-PE (0.20 μg) gMFIs of SPL BCs and ASCs washed 1 or 2 times at 600g or 300g with Foxp3 B2 following fixation. **(G)** Ratio of SPL ASC / BC rIgG2a-PE (0.20 μg) gMFIs from cells washed 1 or 2 times at 600g or 300g with Foxp3 B2 following fixation. Horizontal dashed line represents a ratio of 1. **(B-G)** Data derived from 2 separate experiments. **(C-D, F-G)** Symbols represent individual mice. Horizontal lines represent mean ± SEM.

When we varied the number of post-fixation washes with Foxp3 Buffer 2 (B2) (**Figure 3A**, **red**), we again observed differences in retention. Not surprisingly, BCs showed a reduction in background PE staining which was most evident in the step from 1 to 2 washes (**Figures 3B-C**). Unlike with PFA+Sap, increasing the number of Foxp3 B2 washes did not dramatically enhance ASC retention of PE (**Figures 3B-C**). Rather, the use of multiple washes lowered the overall PE gMFI in ASCs (**Figures 3B-C**) as demonstrated by a left shift of the primary peak of fluorescence intensity. However, the use of 2 or 3 washes induced a separate artifact of PE retention as indicated by the “tails” present in the rIgG2a-PE histograms of multi-washed ASCs (**Figure 3B**). Overall, we did not observe major deviation in the ASC / BC PE gMFI ratio when different numbers of washes were compared (**Figure 3C**). To further characterize this phenomenon, we repeated experiments from **Figure 2** in which we varied post-fixation wash number and RCF. Similar to PFA+Sap, altering RCF did not change PE retention by ASCs (**Figures 3E-G**). While the artifacts induced by number of washes were specific to buffer sets, they shared the common theme of modulation by the number of post-fixation washes.

### 3.4. Fixation and permeabilization method does not differentially impact the analysis of live versus dead antibody-secreting cells

The above experiments indicated that different fixation and permeabilization methodologies induced distinct types of PE retention artifacts in ASCs. Since both protocols are known to impact cellular morphology in their own unique fashion, it is possible that this led to false comparison of ASCs between the 2 methods. That is, a particular method may have included a significant proportion of dead ASCs which may have skewed the results. To exclude this possibility, we repeated the single post-fixation wash PFA+Sap and Foxp3 ICFC protocols with the inclusion of a fixable Live-Dead stain (**Figures 4**). As expected, total ungated events demonstrated a clear positive population indicative of dead cells regardless of the buffer system used (**Figures 4A-B**). When using PFA+Sap, the percentages of gated BCs and ASCs that were Live-Dead^+^ were ~1.8% and 1.0%, respectively (**Figures 4A and 4C**). Interestingly, the use of the Foxp3 buffer set increased the percentage of dead BCs that were analyzed to ~17%. However, only about 0.8% of ASCs analyzed were considered dead (**Figures 4B-C**). Even though the Foxp3 buffer set increased the percentage of dead BCs that were gated, this did not impact PE gMFIs as total and dead BCs showed similar PE background when stained with rIgG2a-PE (0.2 μg) (**Figure 4D**). Overall, these data indicated that different fixation and permeabilization methods had little impact on the analysis of gated, viable ASCs.

**Figure 4:**
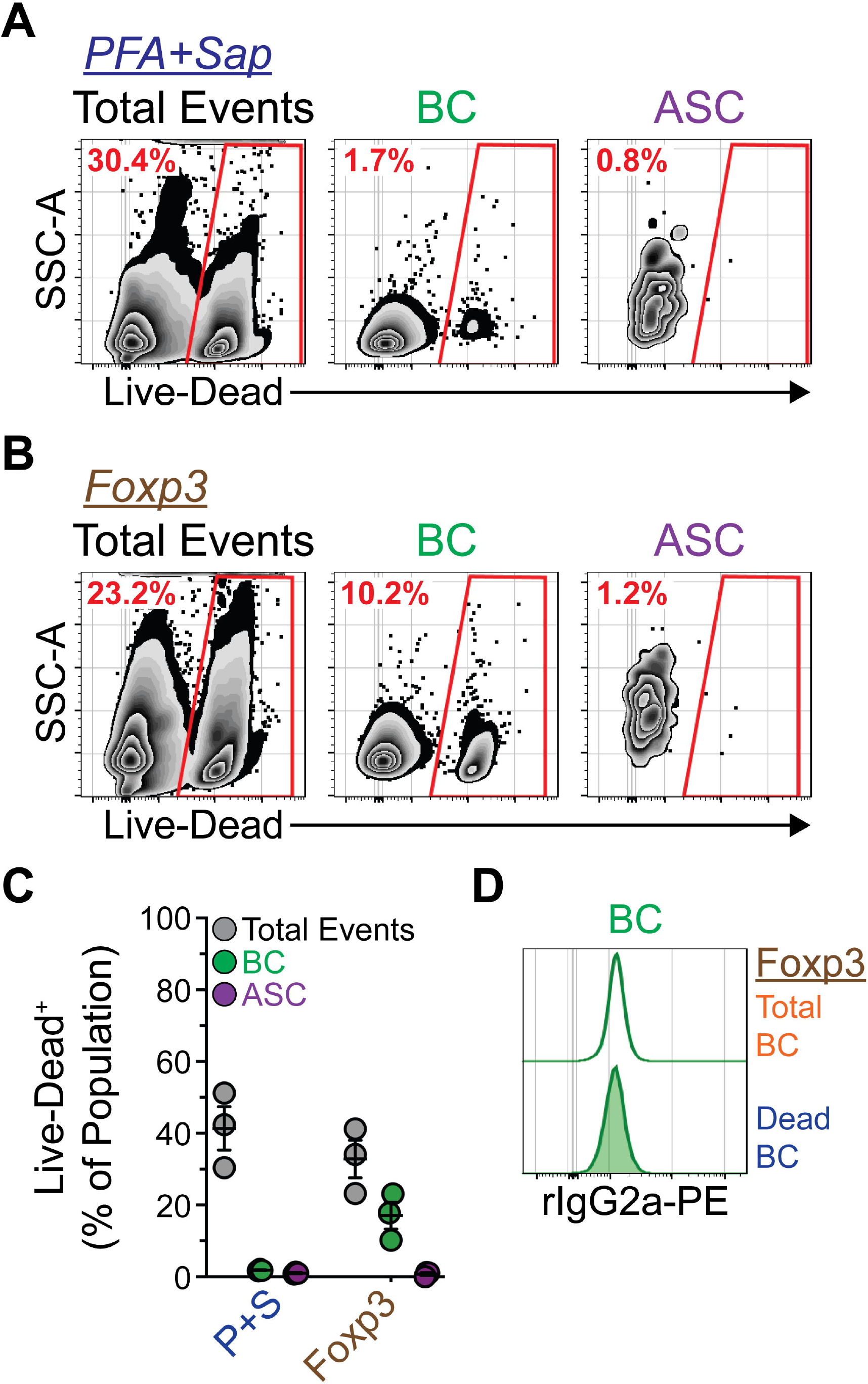
Fixation and permeabilization method does not differentially impact the analysis of live versus dead ASCs. **(A-B)** Representative flow cytometry plots demonstrating Live-Dead staining of total SPL events as well as gated BCs and ASCs. Samples were processed using **(A)** PFA+Sap or the **(B)** Foxp3 buffer set. Dead cells are gated and percentages of Live-Dead^+^ cells are indicated in red. **(C)** Percentages of Live-Dead^+^ cells within total SPL events, BCs and ASCs. Symbols represent individual mice. Horizontal lines represent mean ± SEM. **(D)** Flow cytometry overlay histograms showing rIgG2a-PE fluorescence intensity for total and dead SPL BCs using the Foxp3 buffer set. **(AD)** Data derived from 2 separate experiments.

### 3.5. Performance of one post-fixation wash using the eBioscience Foxp3/Transcription Factor Staining Buffer Set allows for robust intracellular staining of antibody-secreting cells

The above results indicated that a single post-fixation wash protocol using either PFA+Sap or the Foxp3 buffer set may be suitable for ICFC of ASCs when PE-conjugated Abs were present. However, the Foxp3 buffer set, usually in the context of 2 post-fixation washes, is believed to provide increased ICFC utility as it is designed to provide optimal staining of organelle-localized antigens including those in the nucleus (Law et al., 2009). To address these concepts, we compared the use of 1 or 2 post-fixation washes with both buffer sets to detect Pro-IL-1 β and TLR7 (**Figures 5A-D**).

**Figure 5.**
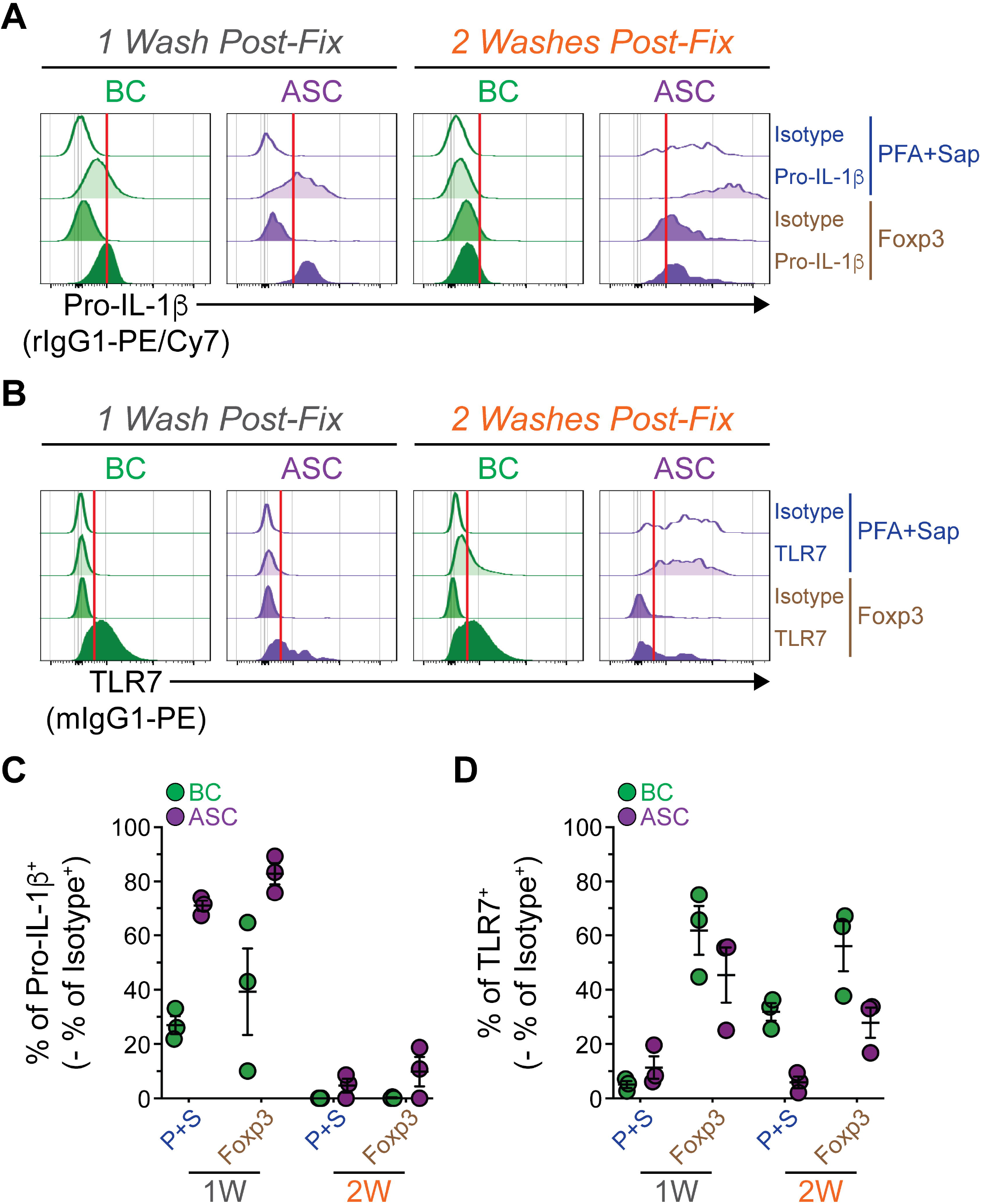
Performance of one post-fixation wash using the eBioscience Foxp3/Transcription Factor Staining Buffer Set allows for robust intracellular staining of ASC proteins. **(A)** Flow cytometry overlay histograms showing rIgG1-PE/Cy7 isotype control and Pro-IL-1 β-PE/Cy7 fluorescence intensity for SPL CD19^+^ BCs, ASCs, PBs and PCs with 1 or 2 post-fixation washes using either PFA+Sap or Foxp3 protocols. **(B)** Flow cytometry overlay histograms showing mouse IgG1 (mIgG1)-PE isotype control and TLR7-PE fluorescence intensity for SPL CD19^+^ BCs, ASCs, PBs and PCs with 1 or 2 post-fixation washes using either PFA+Sap or Foxp3 protocols. **(C)** Percentages of Pro-IL-1β^+^ cells within CD19^+^ BCs and ASCs. Symbols represent individual mice. Horizontal lines represent mean ± SEM. **(D)** Percentages of TLR7^+^ cells within CD19^+^ BCs and ASCs. Symbols represent individual mice. Horizontal lines represent mean ± SEM. **(A-B)** Red vertical lines added to histograms to depict positive staining for Pro-IL-1 β and TLR7. **(A-D)** Data representative of 3 individual experiments.

Pro-IL-1β was chosen as the Immunological Genome Project (ImmGen, https://www.immgen.org) datasets have shown the *Il1b* gene to be preferentially expressed in SPL ASCs, in particular PBs, when compared to total SPL BCs (**Supplementary Figure 2A**). Pro-IL-1β has also been detected in protein lysates from human peripheral blood CD19^+^ B cells (Lim et al., 2020). Finally, cleavage of Pro-IL-1β to its mature IL-1β form and subsequent secretion typically requires integration of multiple “danger” signals (Gurung et al., 2015). As a result, we hypothesized that Pro-IL-1β would accumulate and not be processed under homeostatic conditions thus facilitating its detection. TLR7 was selected as it predominantly localizes to the endosome (Kuznik et al., 2011) allowing for the comparison of cytoplasmic (i.e. Pro-IL-1β) versus organelle-localized staining efficacy. Additionally, *Tlr7* has been shown to be robustly expressed in mouse SPL BCs and ASCs (ImmGen, **Supplementary Figure 2B**) as well as in human tonsillar counterparts (Dorner et al., 2009).

For these experiments, we further gated SPL BCs as CD19^+^ to confirm their identity (**Supplementary Figure 2C**). Fixable Live-Dead staining was also included in all samples to ensure analysis of viable cells. The use of 1 post-fixation wash (**Figure 5A**) allowed for clear detection of Pro-IL-1β with ASCs having an increased percentage of Pro-IL-1b^+^ cells relative to BCs (**Figure 5C**). The use of the Foxp3 buffer set led to modestly increased detection of ASC Pro-IL-1β when compared to PFA+Sap (**Figures 5A and 5C**). In contrast, the use of 2 post-fixation washes resulted in PE/Cy7 retention that prohibited accurate quantification of Pro-IL-1β in ASCs (**Figures 5A and 5C and Supplementary Figure 2D**). Interestingly, detection of Pro-IL-1β was lost in 2 post-fixation wash SPL BCs regardless of buffer set (**Figures 5A and 5C**).

Unlike Pro-IL-1β, TLR7 was minimally detectable when the 1 wash PFA+Sap protocol was utilized (**Figure 5B**). Conversely, the 1 wash Foxp3 protocol resulted in the detection of high amounts of TLR7 (**Figures 5B and 5D**). PFA+Sap in combination with 2 post-fixation washes showed an intermediate TLR7 staining in BCs when compared to results observed with a single post-fixation wash with PFA+Sap and Foxp3 buffers (**Figures 5B and 5D**). Respective to BCs, the 2 wash Foxp3 buffer protocol appeared to be equivalent to the 1 wash version as both displayed ~56-62% average TLR7 positivity with similar gMFIs (**Figures 5B and 5D and Supplementary Figure 2E**). Examination of ASCs again demonstrated PE retention when 2 post-fixation washes were used in the context of PFA+Sap (**Figure 5B**). This was indicated by increased PE gMFIs in isotype control samples (**Figure 5B and Supplementary Figure 2E**). We also observed ASC retention of PE in 2 post-fixation wash Foxp3 buffer set samples (**Figure 5B**). Rather than a dramatic shift in gMFI, the retention observed resembled the previously mentioned “tail” pattern (**Figure 5B**) and resulted in an under estimation of the overall percentage of TLR7^+^ ASCs (**Figure 5D**). In general, both 1 post-fixation wash protocols suppressed PE/Cy7 and PE retention of ASCs for the protein targets shown here. However, the 1 post-fixation wash Foxp3 buffer set protocol provided the most reproducible results and allowed for the detection of cytoplasmic as well as organelle-localized proteins.

## 4. Discussion

While it has been reported that ASCs non-specifically retain PE in the context of ICFC (Bohacova et al., 2021; Kim and Kim, 2013), our current study confirms and extends these findings to include PE/Cy7. A key finding shown here is that the mere act of repetitive centrifugation post-fixation appears to induce the ASC retention of PE. As such, reduction in the number of post-fixation washes and centrifugation steps that occur before ICFC staining ameliorates this phenotype thus allowing for the utilization of extremely bright PE and PE/Cy7 fluorochromes in ASC ICFC.

Previous attempts (Bohacova et al., 2021; Kim and Kim, 2013) to eliminate ASC retention of PE with multiple types of commercial ICFC buffer sets have been unsuccessful. These have included the BD Biosciences CytoFix/Cytoperm buffer set (Kim and Kim, 2013) as well as Foxp3 buffer sets from BD Biosciences, BioLegend and eBioscience/Thermo Fisher Scientific (Bohacova et al., 2021). Interestingly, standard protocols for all of these buffer sets call for multiple centrifugation steps following initial cellular fixation. The only exception to this is the eBioscience Foxp3 buffer set which provides multiple ICFC protocols. One of these, Protocol B, uses a single mandatory centrifugation step post-fixation and was tested and shown to be “curative” in this study (**Figures 3 and 5**). Importantly, a single post-fixation wash also was sufficient to reduce PE retention in the context of PFA+Sap providing some flexibility in what buffers are utilized (**Figures 3 and 5**). This may be an important consideration as different antigens display altered sensitivities in regards to how fixation impacts their Ab-binding epitopes. In regard to the protein targets tested here, optimal staining of ASC intracellular proteins was observed while using the eBioscience Foxp3/Transcription Factor Staining Buffer Set rather than PFA+Sap (**Figure 5**). This was most obvious when 1 post-fixation wash was performed (**Figure 5**) and may simply be a result of the permeabilization agents that differ between protocols. Sap is a “weak” detergent which has been used to permeabilize fixed cells for ICFC; however, instances have been observed in which Sap does not effectively permeabilize certain types of membranes (Mercanti and Cosson, 2010). Under these circumstances, Triton X-100 has served as a more appropriate method as this “strong” detergent is commonly used for immunofluorescence of nuclear proteins (Spector, 2011). While the permeabilizing agent contained within the eBioscience Foxp3/Transcription Factor Staining Buffer Set used here is not explicitly stated, similar kits such as the Human FoxP3 Buffer Set (Catalog # 560098) from BD Biosciences is known to contain methanol, a common reagent used to fix and permeabilize cells during ICFC (Levitt and King, 1987). Relatedly, we did observe differences in the type of PE retention when PFA+Sap and Foxp3 buffer sets were compared. We speculate that this may simply be a consequence of how the different buffer sets interact with cells. For example, PFA fixation leads to protein-protein crosslinks that also preserve overall cell structure. In contrast, methanol is an organic solvent which precipitates proteins through dehydration resulting in distortion of cellular morphology. This is readily observed when forward and side scatter properties of cells treated with these reagents are examined on a flow cytometer. However, the idea remains to be tested. Lastly, we did find some variance between isotypes in the magnitude of PE and PE/Cy7 retention (**Figure 1 and Figure 3 versus Figure 5**). While we do not currently know the cause of this phenomenon, it may be related to differences in constant domain structure between antibody isotypes (Schroeder and Cavacini, 2010). For example, differences in lysine amino acid number may directly impact efficiency of fluorochrome conjugation (Nanna et al., 2017). In this sense, increased lysine residues within a particular isotype could lead to a higher ratio of fluorochrome conjugation and thus higher retention of PE.

An obvious question becomes why do ASCs exhibit PE retention and by extension, why does centrifugation augment this retention phenotype? While our study does not directly address this, a possibility may be related to the major function of ASCs which is the production of Abs. Of note, Abs are heavily glycosylated (Irvine and Alter, 2020) and various compounds exist that bind carbohydrates (Sun et al., 2016) facilitating their detection independent of specific antibody:antigen interactions. Therefore, it is tempting to speculate that the high abundance of intracellular immunoglobulins combined with repetitive centrifugation can induce PE retention through incidental damage to cellular glycoproteins resulting in exposure of PE-interacting surfaces or structures. Alternatively, ASCs possess an expanded endoplasmic reticulum (ER) and Golgi network which may also suffer from the same post-centrifugation consequences hypothesized for Abs. Along these lines, it would be of interest to investigate PE retention in other cells which possess an expanded ER compartment such as plasmacytoid dendritic cells (pDCs) (Colonna et al., 2004). Would the deletion of *Xbp1* which leads to loss of the expanded ER network in ASCs (Taubenheim et al., 2012) and pDCs (Iwakoshi et al., 2007) result in the loss of PE retention?

Regardless of causation, the ability to ameliorate PE retention, as shown here, now facilitates the usage of PE- and PE/Cy7-conjugated Abs in assessing intracellular protein expression by ASCs. This is relevant to stand alone analysis of ASCs as well as comparison to other cell types such as naive or activated BCs. This is a critical point as BC to ASC differentiation has been correlated with acquisition of selected cytokine expression (Shi et al., 2015; Suzuki-Yamazaki et al., 2017) and could theoretically serve as a key diagnostic tool in the right context (Fink, 2012). As such, reducing any potential for a false positive is critical (Savasan et al., 2018). To this end, FMOs and isotype control Abs have both been utilized in the field of flow cytometry and hotly debated in regards to what the “correct” strategy is. We would suggest that much like anything else, this depends on context. For ICFC of ASCs, the use of PE- and PE/Cy7-conjugated isotype control Abs is absolutely essential in identifying a retention phenotype (**Figure 1**) (Bohacova et al., 2021; Kim and Kim, 2013) that would have been misinterpreted for positive staining if only FMO samples and those stained with target specific Abs were assessed.

Finally, we provide biological insight at the protein level into the evolution of ASCs and their potential to interact with their surrounding environment. RNA sequencing of mouse bone marrow plasma cells (Pioli et al., 2019) as well as quantitative polymerase chain reaction of human peripheral blood plasmablasts and tonsillar plasma cells (Dorner et al., 2009) has demonstrated expression of *Tlr7* at the gene level. We show here that a portion of SPL ASCs express TLR7 (**Figure 5**). Given the role of this protein during viral responses (Lester and Li, 2014), it will be interesting to see how ASC expression of TLR7 changes during infection and whether or not this contributes to the functionality of these cells. Furthermore, we show increased Pro-IL-1β expression in SPL ASCs compared to their upstream BC counterparts (**Figure 5**). This coincides with RNA sequencing data available in the ImmGen database (http://rstats.immgen.org/Skyline/skyline.html) which illustrates enriched expression of *Il1b* transcripts in SPL plasmablasts relative total BCs (**Supplementary Figure 2A**).

In summary, we have identified a key contributing factor to the ASC retention of PE molecules during ICFC. It should be noted that this and other studies (Bohacova et al., 2021; Kim and Kim, 2013) regarding ASC retention of PE have focused on murine cells. Human ASCs are ultra-structurally similar to those from mice (e.g. expanded ER network); however, it is currently unknown if human ASCs suffer from the same PE retention. Regardless, we provide a framework to not only determine the existence of PE retention in human ASCs and other cell types (e.g. pDCs) but to also ameliorate its impact on ICFC staining.

## Author Contributions

KATP and PDP conceptualized the study. PR, MC, MK and KATP performed experiments. KATP and PDP analyzed data. PDP wrote the manuscript and all authors approved the final version.

## Acknowledgments

Flow cytometry was performed at the WMed Flow Cytometry and Imaging Core which is managed by Michael Clemente.

---

**Supplementary Figure 1: PE- and PE/Cy7-conjugated Abs are retained by ASCs in a concentration dependent manner.** Flow cytometry-derived geometric mean fluorescence intensities (gMFIs) for various isotype control Abs conjugated to **(A-B)** APC, **(C-D)** PE and **(E-F)** PE/Cy7. **(A, C, E)** Data shown for B220^+^ CD138^-^ BCs and B220^+/-^ CD138^HI^ CD90.2^-^ IgD^-^ ASCs using 0.04 and 0.20 μg per Ab as well as FMO controls. **(B, D, F)** Data shown for B220^+^ CD138^HI^ CD90.2^-^ IgD^-^ PBs and B220^-^ CD138^HI^ CD90.2^-^ IgD^-^ PCs using 0.04 and 0.20 μg per Ab as well as FMO controls. **(A-F)** Symbols represent individual mice from 3 separate experiments. Horizontal lines represent mean ± SEM.

**Supplementary Figure 2: One post-fixation wash using the eBioscience Foxp3/Transcription Factor Staining Buffer Set allows for robust intracellular staining of ASC proteins. (A)** ImmGen *Il1b* expression data for SPL BCs, PBs and PCs. **(B)** ImmGen *Tlr7* expression data for SPL BCs, PBs and PCs. **(C)** Representative flow cytometry plots demonstrating CD19 gating within the BC population compared to total singlets using either PFA+Sap or Foxp3 protocols. Red vertical lines added to histograms to depict positive staining for CD19. **(D)** Flow cytometry-derived gMFIs for PE/Cy7-conjugated isotype control and Pro-IL-1 β Abs using 1 post-fixation wash and 2 post-fixation wash versions of the PFA+Sap (P+S) and Foxp3 protocols. **(E)** Flow cytometry-derived gMFIs for PE-conjugated isotype control and TLR7 Abs using 1 post-fixation wash and 2 post-fixation wash versions of the PFA+Sap (P+S) and Foxp3 protocols. **(A-B)** ImmGen data represent expression values normalized by DESeq2. Data were downloaded and replotted as is without further calculation. **(D-E)** Symbols represent individual mice from 3 separate experiments. Horizontal lines represent mean ± SEM.

