## Supplementary figures and images for "Intracellular flow cytometry staining of antibody-secreting cells using phycoerythrin-conjugated antibodies: pitfalls and solutions"

### Supplementary Figure 1

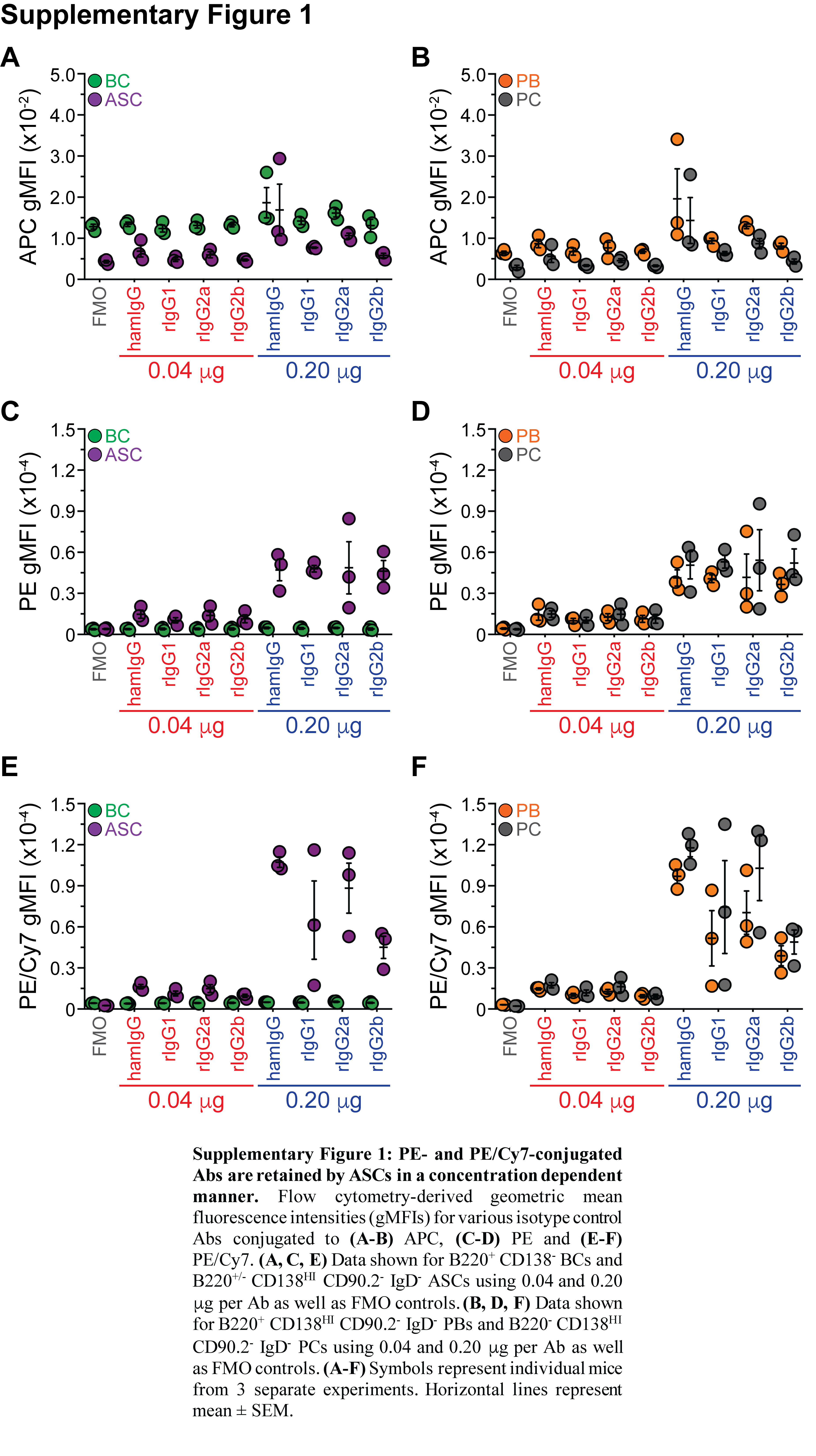

### Supplementary Figure 2

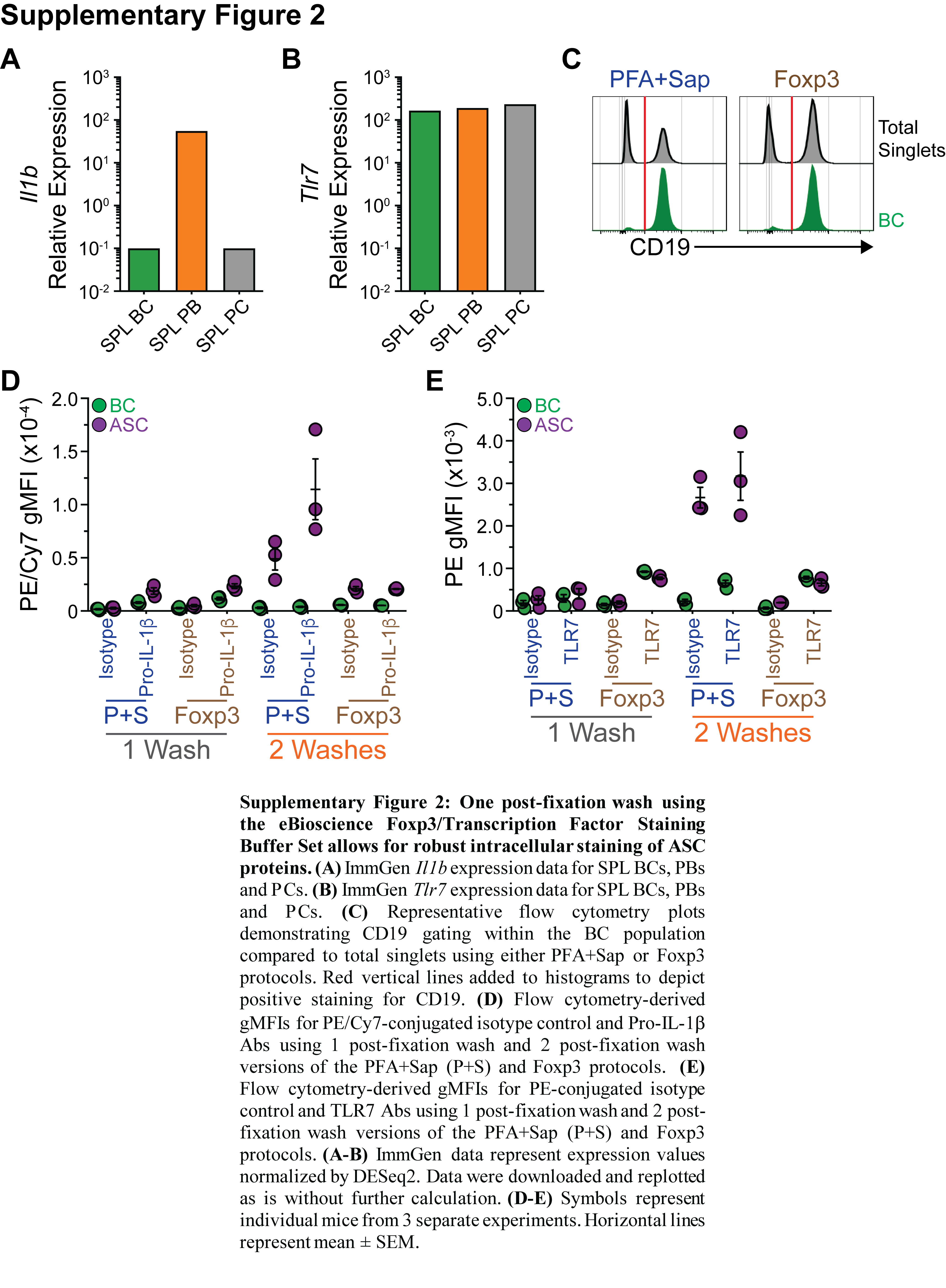
