## Supplementary Table 1 for "Intracellular flow cytometry staining of antibody-secreting cells using phycoerythrin-conjugated antibodies: pitfalls and solutions"

**Supplementary Table 1: List of antibodies used in this study.**

| Antibody | Source | Identifier |
|---|---|---|
| CD19-BUV395 (Clone: 1D3) | BD Biosciences | Cat# 563557; RRID: AB_2722495 |
| CD45R(B220)-PerCP/Cy5.5 (Clone: RA3-6B2) | BD Biosciences | Cat# 552771; RRID: AB_394457 |
| CD45R(B220)-BV711 (Clone: RA3-6B2) | BD Biosciences | Cat# 563892; RRID: AB_2738470 |
| CD138-BV421 (Clone: 281-2) | BD Biosciences | Cat# 562610; RRID: AB_11153126 |
| IgD-BV605 (Clone: 11-26c.2a) | BioLegend | Cat# 405727; RRID: AB_2562887 |
| IgD-APC-H7 (Clone: 11-26c.2a) | BD Biosciences | Cat# 565348; RRID: AB_2739201 |
| CD90.2(Thy-1.2)-BV605 (Clone: 53-2.1) | BD Biosciences | Cat# 563008; RRID: AB_2665477 |
| CD90.2(Thy-1.2)-APC/Cy7 (Clone: 53-2.1) | BD Biosciences | Cat# 561641; RRID: AB_10898013 |
| CD16/32-Unlabeled (Clone: 93) | Thermo Fisher Scientific | Cat# 14-0161-86; RRID: AB_467135 |
| Mouse IgG-Unlabeled | SouthernBiotech | Cat# 0107-01; RRID: AB_2732898 |
| Armenian Hamster IgG-APC Isotype Control (Clone: HTK888) | BioLegend | Cat# 400911 |
| Armenian Hamster IgG-PE Isotype Control (Clone: HTK888) | BioLegend | Cat# 400908 |
| Armenian Hamster IgG-PE/Cy7 Isotype Control (Clone: HTK888) | BioLegend | Cat# 400922 |
| Rat IgG1-APC Isotype Control (Clone: RTK2071) | BioLegend | Cat# 400412 |
| Rat IgG1-PE Isotype Control (Clone: RTK2071) | BioLegend | Cat# 400408 |
| Rat IgG1-PE/Cy7 Isotype Control (Clone: RTK2071) | BioLegend | Cat# 400416 |
| Rat IgG2a-APC Isotype Control (Clone: RTK2758) | BioLegend | Cat# 400512 |
| Rat IgG2a-PE Isotype Control (Clone: RTK2758) | BioLegend | Cat# 400507 |
| Rat IgG2a-PE/Cy7 Isotype Control (Clone: RTK2758) | BioLegend | Cat# 400521 |
| Rat IgG2b-APC Isotype Control (Clone: RTK4530) | BioLegend | Cat# 400612 |
| Rat IgG2b-PE Isotype Control (Clone: RTK4530) | BioLegend | Cat# 400608 |
| Rat IgG2b-PE/Cy7 Isotype Control (Clone: RTK4530) | BioLegend | Cat# 400617 |
| Pro-IL-1β-PE/Cy7 (Clone: NJTEN3) | Thermo Fisher Scientific | Cat# 25-7114-82; RRID: AB_2573526 |

| | | |
|---|---|---|
| Mouse IgG1-PE Isotype Control (Clone: MOPC-21) | BD Biosciences | Cat# 554680 |
| CD287(TLR7)-PE (Clone: A94B10) | BD Biosciences | Cat# 565557; RRID: AB_2739295 |
